# Unraveling the GM_1_ specificity of Galectin-1 binding to lipid membranes

**DOI:** 10.1101/2024.09.20.614102

**Authors:** Federica Scollo, Waldemar Kulig, Gabriele Nicita, Anna-K. Ludwig, Joana C. Ricardo, Valeria Zito, Peter Kapusta, Ilpo Vattulainen, Marek Cebecauer, Hans-Joachim Gabius, Herbert Kaltner, Giuseppe Maccarrone, Martin Hof

## Abstract

Galectin-1 (Gal-1) is a galactose-binding protein involved in various cellular functions. Gal-1’s activity has been suggested to be connected to two molecular concepts, which are however lacking experimental proof: a) enhanced binding affinity of Gal-1 towards membranes containing monosialotetrahexosylganglioside (GM_1_) over disialoganglioside GD_1_a and b) cross-linking of GM_1_’s by homodimers of Gal-1. We provide evidence about the specificity and the nature of Gal-1 interaction with model membranes containing GM_1_ or GD_1_a, employing a broad panel of fluorescence-based and label-free experimental techniques, complemented by atomistic biomolecular simulations. Our study demonstrates that Gal-1 binds indeed specifically to GM_1_, and not to GD_1_a, when embedded in membranes over a wide range of concentrations (i.e., 30 nM to 10 μM). The apparent binding constant is about tens of micromoles. On the other hand, no evidence of Gal-1/GM_1_ cross-linking was observed. Our findings suggest that cross-linking does not result from sole interactions between GM_1_ and Gal-1, indicating that in a physiological context, additional triggers are needed, which shift the GM_1_/Gal-1 equilibria towards the membrane-bound homodimeric Gal-1.

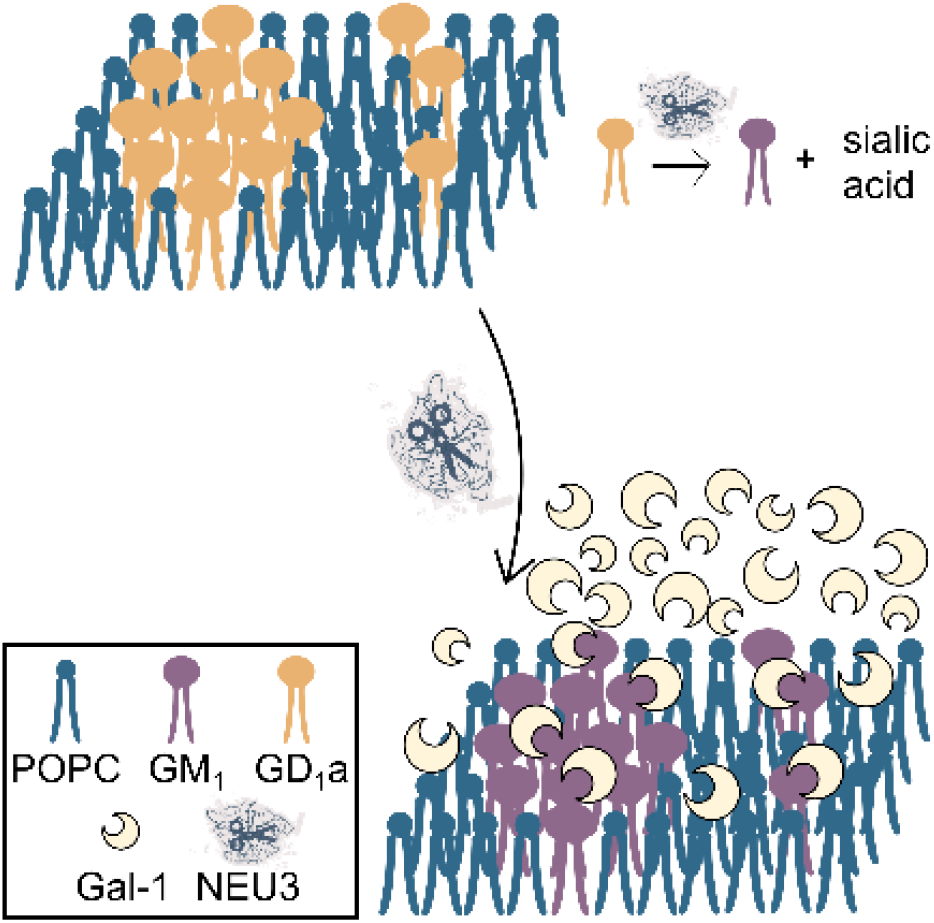

## Introduction

Galectins are a ubiquitous family of galactose-binding proteins in the nucleus, cytosol, and extracellular matrix. ^1^ They are involved in various biological processes, from RNA splicing to cell growth regulation, including cell adhesion, embryogenesis, inflammation and immune function, apoptosis, angiogenesis, and tumor metastasis. ^2-15^ All galectins have a common feature: they possess one (or more) β-sandwich carbohydrate recognition domain (CRD) by which they interact with molecules having a galactoside moiety. ^16^ Sixteen human galectins have been discovered, ^17^ commonly classified into three structurally different types, i.e., prototype, chimera, and tandem-repeat type, based on the structural presentation of their CRD. ^18^ Galectin-1 (Gal-1), which belongs to the prototype class, ^19, 20^ was the first of the galectins to be characterized. Nevertheless, there is still little known about its specific roles *in vivo* and the underlying working mechanisms. ^21^ One of these uncertainties concerns its specific binding to GM_1_, one of the most abundant glycosphingolipids located at the outer layer of the plasma membrane ^22, 23^, which has been postulated to be crucial for a variety of its cellular functions. The intercellular communication of regulatory and effector T cells (T_reg_ and T_eff_, respectively) might serve here as a clinically relevant example, in which the reduction of Gal-1/GM_1_ interactions inevitably leads to imbalances in T cell communication and may causally trigger autoimmune diseases. ^24^ Antigen-presenting cells (APC) activate T cell receptors of T_reg_ cells, causing upregulation of Gal-1 and its release from T_reg_ cells into the surroundings. Simultaneous activation of T_eff_ cells by APC causes increased formation of GM_1_ through enhanced neuraminidase 3 activity, converting disialoganglioside GD_1_a to GM_1_ by removing the terminal sialic acid moiety. It has been suggested that the upregulated Gal-1 forms homodimers and cross-links to GM_1_, leading to co-cross-linking with GM_1_-associated heterodimeric integrin. That cross-linking might lead to initiating a post-binding signaling cascade that activates a TRPC5 Ca^2+^ channel, which in turn blocks T_eff_ cell proliferation. ^8, 9, 11, 24-26^

This example demonstrates how Gal-1 translates the metabolic conversion of ganglioside GD_1_a to GM_1_ by neuraminidase into a specific response on the cellular level. ^27^ Considering the crucial role of the specific Gal-1/GM_1_ interaction, ^28^ it is essential to experimentally prove that specificity. The strongest evidence for that specificity comes from experiments in neuroblastoma cells. ^19, 27, 29^ In these contributions, the Gal-1 binding to the cell surface is quantified by radioisotope I^125^ Gal-1 marking ^19, 29^ or by Gal-1 visualization via fluorescently labelled antibodies. ^27^ The concept of these experiments is a) to vary GM_1_ concentrations on the cell surface by using different cell lines or interfering with the neuraminidase activity or b) blocking surface GM_1_ by antibodies or cholera toxin B subunit. These experiments indicate that the surface concentration of Gal-1 is correlated with the availability of the GM_1_ pentasaccharide headgroup. Nevertheless, these *in vivo* experiments do not give a direct proof for Gal-1 binding to GM_1_, which gives the motivation to characterize the Gal-1/GM_1_ interaction *in vitro*.

Frontal affinity chromatography or NMR experiments have been performed with the isolated GM_1_ pentasaccharide. ^30, 31^ As the lipid moiety was neglected, these experiments show that the GM_1_ pentasaccharide headgroup with its terminal galactose moiety provides a docking site for Gal-1, but not that Gal-1 binds to GM_1_ when embedded in a membrane environment. To the best of our knowledge, an X-ray reflectivity and grazing incidence diffraction study on a lipid monolayer at the air/water interface ^32^ and a chemically induced dynamic nuclear polarization study using GM_1_ in dodecylphosphocholine micelles ^30, 31^ are the only *in vitro* studies using GM_1_. Of note, significant efforts were made using supramolecular self-assembled polymers with controlled size and high glycans surface concentration to investigate galectin avidity. ^33, 34^ However, these studies do not provide insight into the GM_1_ specificity since they are based on lactose glycodendrimers. Altogether, these *in vitro* studies do not yield a convincing proof for a specific Gal- 1/GM_1_ interaction.

Moreover, it has been suggested that GM_1_ cross-linking by homodimer Gal-1 might lead to 2-dimensional glycan-galectin aggregates, called Gal-1 “lattices”. ^15, 28, 35, 36^ Gal-1 and Gal-1 “lattice” are claimed to play a crucial role in glycocalyx organization and regulation. ^37^ However, the literature does not provide any direct proof for GM_1_ being involved in Gal-1 cross-linking or surface aggregation. The apparent imbalance between the suggested cellular role of the Gal-1/GM_1_ interaction and the lack of experimental data calls for a thorough characterization of Gal-1 interactions with GM_1_ embedded in well-controlled membrane model systems.

In this work, we investigate the interaction of wild type Gal-1 with model membranes containing GM_1_ or GD_1_a, combining fluorescence based and label-free techniques. Specifically, we employed Förster resonance energy transfer (FRET) approach, confocal fluorescence Microscopy (FM), quartz crystal microbalance with dissipation monitoring (QCM-D), and isothermal titration calorimetry (ITC). The experiments are complemented by all-atom molecular dynamics (MD) simulations to gain insights into the Gal-1/GM_1_ specificity at a molecular level. All these techniques provide clear evidence for the GM_1_ specificity of Gal-1 binding to lipid membranes characterized by micromolar binding constants.

## Results and Discussion

On a cell surface, glycosphingolipids are prominently encountered and tightly controlled. Shifts in ganglioside profiles occur in cell activation/differentiation or neuronal regeneration. The conversion of ganglioside GD_1_a to GM_1_ via enzymatic desialylation by the plasma membrane neuraminidase 3 has been suggested to be read by a concerted upregulation of Gal-1, with the ensuing growth control of neuroblastoma cells, axon regeneration and effector/regulatory T cell communication in autoimmunity control. ^28^ This fascinating concept is based on the elevated binding affinity of Gal-1 towards GM_1_ over GD_1_a when embedded in membranes, an assumption which, however, lacks experimental proof.

To characterize lipid specificity of Gal-1 binding, phospholipid bilayers are an ideal model system. More specifically, we monitored Gal-1 binding to diverse unilamellar vesicles composed of palmitoyl-oleoyl phosphatidylcholine (POPC) with and without GM_1_ or GD_1_a in relevant concentrations of 4 mol%, the highest reported levels of GM_1_ in neurons. ^28, 38, 39^ At these ganglioside concentrations, GM_1_ and GD_1_a organize into transient and fluid nanodomains. ^40, 41^ First, we used unlabeled Giant Unilamellar Vesicles (GUVs) containing POPC with 4 mol% of GM_1_ to visualize their interaction with Gal-1 labelled with tetramethylrhodamine dye (TMR), referred to here as Gal-1/TMR (for detailed characterization see Supplementary Information (**SI**)). The GUVs were visualized using confocal microscopy. Single-channel detection revealed a fluorescence emission on the GUVs in the spectral emission range of the TMR, indicating that the Gal-1/TMR localizes and accumulates on the POPC:GM_1_ GUVs, unlike on the POPC or POPC:GD_1_a GUVs (as shown in **Fig. 1a**). Fluorescence Microscopy data are summarized in **Fig. 1b** and show that Gal-1/TMR adsorbs onto (84 ± 6) % of POPC:GM_1_ GUVs membranes and only onto (15 ± 5) % of POPC GUVs and (11 ± 6) % of POPC:GD_1_a GUVs, suggesting a specific interaction between Gal-1 and GM_1_.

**Figure 1.**
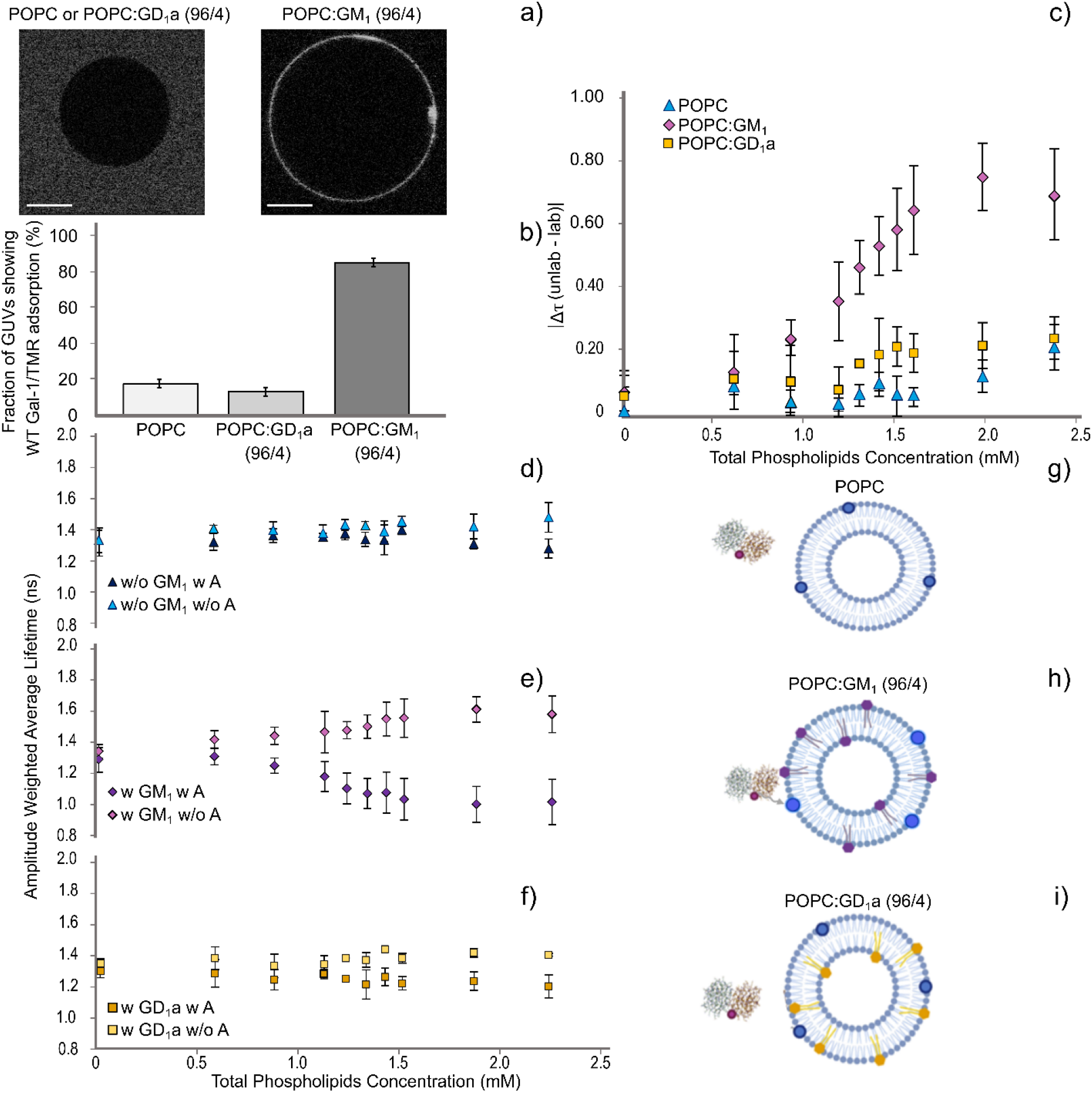
Representative confocal images of Gal-1/TMR with unlabeled GUVs and FRET data measured on Gal-1/TMR and labelled and unlabeled LUVs. a) Representative xy cross-sections of unlabeled POPC or POPC:GD_1_a and POPC:GM_1_ incubated with Gal- 1/TMR. Scale bars are shown on the bottom left (10 μm). b) Fraction of GUVs showing adsorption and associated error of Gal-1/TMR at the membrane of POPC, POPC:GD_1_a and POPC:GM_1_ (see **Table S1** for the details). n > 15 (at least three different electroformations). c) |Δτ (unlab - lab)| as a function of total phospholipids concentration for three different compositions, i.e., POPC (blue triangles), POPC:GM_1_ (purple rhombs), and POPC:GD_1_a (yellow squares). |Δτ (unlab - lab)| is defined as the absolute value of the differences between the amplitude weighted average lifetime of the Gal-1/TMR (the donor) in the presence of LUVs non-containing the acceptor (labelled as A) and the same in the presence of LUVs containing the acceptor (1% of DOPE-Atto 633). Error bars represent the S.E.M. d, e and f) Amplitude weighted average lifetime of the Gal-1/TMR as a function of increasing dispersion concentration of Gal-1/TMR and POPC, POPC:GM_1_ and POPC:GD_1_a, respectively. All the decay curves were acquired at 25°C. Error bars represent the SD (n=5 for POPC and POPC:GM_1_ and n=2 for POPC:GD_1_a). The average was calculated on samples prepared by at least two different extrusions. g, h, and i) Schematic cartoon of Gal-1/TMR and POPC, POPC:GM_1_ and POPC:GD_1_a LUVs, respectively. The structure of the protein was taken from PDB (https://www.rcsb.org/3d-view/1SLA). TMR and DOPE-Atto 633 dyes are shown in the cartoon as blue and red dots, respectively. POPC, GM_1_, and GD_1_a were done in Biorender Software and represented as light blue, purple, and yellow, respectively. The grey arrow shows the occurrence of the energy transfer from the donor (Gal-1/TMR) to the acceptor (POPC:GM_1_ + DOPE-Atto 633).

To better decipher this specificity, we performed FRET experiments. Herein, we focused on the fluorescence lifetime of the donor (Gal-1/TMR), and monitored lifetime changes upon titration of Large Unilamellar Vesicles (LUVs) composed of POPC, POPC:GM_1_, or POPC:GD_1_a into the galectin-containing solution, using 1% of DOPE-Atto 633 in these lipid mixtures as a FRET acceptor. **Fig. 1g, h**, and **i** illustrate cartoon representations of the systems used. These data show that the lifetime of the protein remains constant when adding labelled and unlabeled POPC as well as POPC:GD_1_a (**Fig. 1d, f**). However, the lifetime of Gal-1/TMR in the presence of POPC:GM_1_ changes significantly (**Fig. 1e**). Specifically, titrating with LUVs embedded with the acceptor causes a decrease of the donor lifetime, thus indicating the occurrence of FRET (**Fig. 1e**).

Conversely, we observed an increase in the TMR fluorescence lifetime in the absence of the acceptor (**Fig. 1e**). This can be explained by different nano-environments sensed by the dye linked to Gal-1 due to the change from bulk to the membrane-bound state. For a better visualization of the data, we plotted the absolute difference in Gal-1/TMR lifetime in the absence and in the presence of the acceptor-containing vesicles against the total concentration of phospholipids (Δτ (unlab - lab) – **Fig. 1c**). These data (**Fig. 1c, d, e**, and **f**) show that the presence of GM_1_ in the bilayer affects the WT Gal-1/TMR’s lifetime. This effect is specific to GM_1_, and not to its precursor GD_1_a.

The titration curves in **Fig. 1e** show a threshold lipid concentration of higher than 0.6 mM for a detectable change in the fluorescence time. Apparently, the binding of fluorescently labelled Gal-1 is only detectable at such elevated lipid concentrations. As shown in the **SI** (**Fig.S3b**), UV-Vis characterization of the WT Gal- 1/TMR indicates that about 55% of the protein is unlabeled and 45% of the Gal-1 molecules are labelled with one or more fluorophores. We suggest that unlabelled Gal-1 predominately binds at lipid concentrations lower than 0.6 mM, which trivially cannot be detected by fluorescence. Thus, the apparent threshold lipid concentration of 0.6 mM for the change of the fluorescence lifetime is a consequence of a significant decrease of Gal-1 binding affinity due to TMR labelling. It is indeed generally known that labelling might decrease binding affinities of proteins. ^42-44^. In summary, fluorescence experiments demonstrate the specificity of the Gal-1 interaction with GM_1_ or GD1a containing membranes. However, the covalent attachment of the TMR fluorescent dye apparently decreases the binding affinity, a finding which should be kept in mind when using fluorescence-based techniques when characterising the binding of galectins to membranes.

Next, we quantified the binding of unlabeled Gal-1 to LUVs using label-free techniques, specifically QCM-D and ITC. QCM-D is a surface technique that quantifies the material deposited onto the sensor with a sensitivity of the order of ng/cm^2. 45^ In each experiment, we monitored frequency changes induced by the deposition of intact LUVs (**Fig. 2b**, step I – see the **S.I**. for the details) and after the injection of the protein (**Fig. 2b**, step III). The presence of 4 mol% of GM_1_ or GD_1_a in the vesicles introduces negative charges, which are not present in the POPC vesicles. This leads to more significant electrostatic repulsive interactions between the adsorbed vesicles and prevents the complete coverage of the gold sensor. To ensure consistent coverage of the sensor in experiments with different lipid compositions, a second step of surface coating with uncharged POPC lipid vesicles (**Fig. 2b**, step II) was introduced (see **S.I**., **Table S3** and **Fig. 2b** for a schematic representation). A representative frequency plot for each lipid composition is presented in **Fig 2a**. To gain insight into the binding of Gal-1 to lipid membranes, we focused on the frequency shifts due to the injection of 10 μM Gal-1 (**Fig. 2a** and **b**, step III). This frequency is obtained by calculating the difference between the final frequency reached after the washing step and the initial frequency that was observed prior to the addition of the protein (see **S.I**. for details). The results are presented in **Fig. 2c** as a bar plot. These experiments yielded the following observations: a) a small increase in frequency was noted for POPC (Δf = - 6 Hz ± 2 Hz), probably attributed to the slow detachment of vesicles from the sensor; b) Gal-1 did not cause a significant change in frequency to the previously deposited layer of POPC:GD_1_a vesicles; c) intriguingly, the addition of Gal-1 to POPC:GM_1_ LUVs led to a significant decrease in the frequency (Δf = 25 Hz ± 2 Hz), providing evidence that Gal-1 binds to the POPC:GM_1_ LUVs, consistent with findings from of fluorescence-based experiments (**Fig. 1**). The fraction of protein bound was calculated by dividing each frequency shift related to a certain concentration of the Gal-1 to the highest frequency shift observed (i.e., 25 Hz at 10 μM of Gal-1). The results are reported in **Fig. 2d**, illustrating the fraction bound as a function of different concentrations of Gal-1. The curve was then fitted, yielding an apparent K_d_ of (4 ± 1) μM. Of note, we deposited intact POPC LUVs containing physiologically relevant concentrations of GM_1_ instead of forming Supported Lipid Bilayers (SLBs) and thus avoided the influence of the support on the physical-chemical properties of the bilayer. The apparent K_d_ is in the same range as estimated by radioisotope I^125^ Gal-1 marking experiments to neuroblastoma cells. ^19, 27, 29^

**Figure 2.**
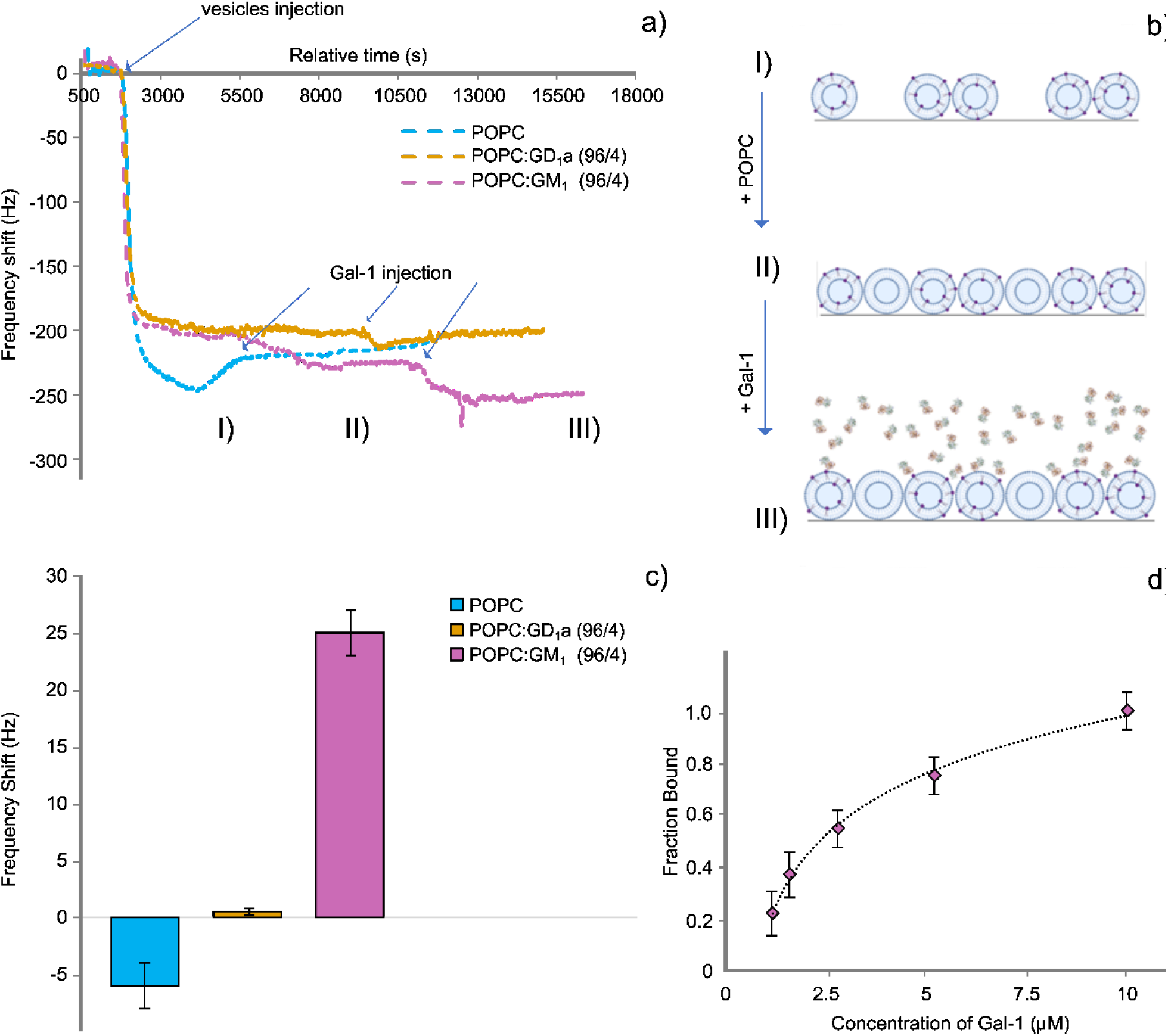
QCM-D data. a) Representative frequency plots for POPC, POPC:GM_1_ and POPC:GD_1_a vesicles deposition on the QCM-D sensor, followed by injecting additional POPC (if needed) and Gal-1. b) Cartoon representing the three steps of the experiment flow. I) injection of the lipid vesicle of interest II) injection of additional POPC in case of GM_1_/GD_1_a containing vesicles and III) injection of the Gal-1. c) Δf of the III step calculated as frequency after – before Gal-1 injection for three POPC (blue bar), POPC:GM1 (purple bar) and POPC:GD_1_a (yellow bar). Error bars signify the S.E.M (n=3 for POPC:GM_1_ and n=2 for POPC and POPC:GD_1_a) d). Fraction of the Gal-1 to POPC:GM1 vesicles as a function of Gal-1 concentration. The fraction bound was calculated by dividing each frequency shift by the maximum frequency shift achieved at the saturation point, thus the maximum amount of protein that can bind to the supported lipid vesicles. The same experiment flow was adopted in a) and d) (described in b)) but varying the concentration of the Gal-1 at the third step. For each concentration, the experiment was repeated in triplicates. Error bars represent the S.E.M (n=3). All the data were acquired at 25 – 30 °C.

Nevertheless, K_d_s determined by surface-based methods as QCM-D can deviate from solution methods for several reasons, i.e., chemical heterogeneity due to the immobilization procedure, crowding/steric hindrance, or restriction in the degree of freedom or diffusional properties. ^46, 47^ As this can modify the thermodynamics of the interaction, we employed ITC to gain detailed insight into the thermodynamics of Gal-1/GM_1_ interaction under equilibrium conditions without vesicle immobilization. ^48^ Titrating the Gal-1 protein in a buffer with the dispersion of POPC:GM_1_ LUVs enabled us to determine the heat released or absorbed by the protein-lipid membrane interaction (**Fig. 3a**).

**Figure 3.**
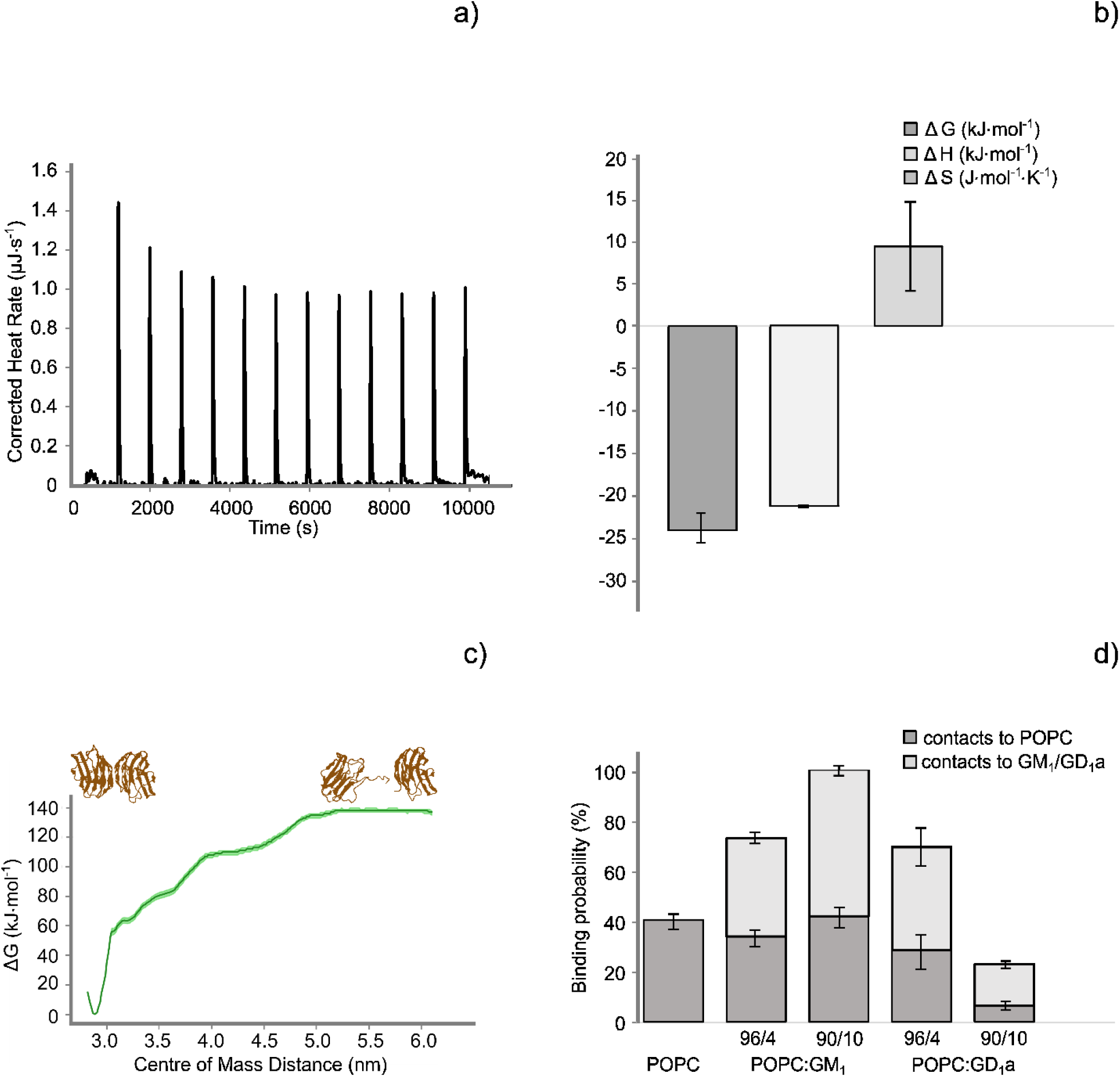
Thermodynamic study: Isothermal Titration Calorimetry and all-atom MD simulations. a) Representative thermogram depicting the corrected heat rate measured when adding POPC:GM_1_ (96/4) to Gal-1 as a function of time. Controlled and precise additions of POPC:GM_1_ LUVs were performed to a solution of Gal-1 (10 mM PBS, 137 mM NaCl, 0.27 mM KCl, pH=7.4). Keeping the same instrumental setup, analogous additions to PBS (without Gal-1) of POPC:GM_1_ LUVs were made. These control experiments were necessary to obtain the blank titrations, which were subtracted from the sample measurements in the context of analyzing the reaction heats. b) Thermodynamic parameters (ΔG, ΔH, ΔS) and their associated error bars indicating the standard deviation obtained titrating POPC:GM_1_ (96/4) to a Gal-1 solution. The thermodynamic data were gained from a global analysis performed by using HypCal software (n=3). c) Potential of mean force (PMF) of Gal-1 dimer dissociation in water obtained using the umbrella sampling techniques. The PMF was calculated using the weighted histogram analysis method implemented in GROMACS. Errors bands represent the errors calculated using the Bayesian bootstrapping method. d) Probability of the Gal-1 dimer binding to five different bilayers, i.e., POPC, POPC:GM_1_ (96/4), POPC:GM_1_ (90/10), POPC:GD_1_a (96/4), POPC:GD_1_a (90/10), represented by the different bars. The bars’ dark and pale grey portions represent the contacts to POPC and GM_1_ or GD_1_a in each different bilayer composition, respectively (see **Table S2** for details).

We employed HypCal software ^49^ to determine thermodynamic parameters of the binding. Three curves were globally analyzed. We found an exothermic reaction with a related ΔH of (-21.39 ± 0.15) kJ mol^-1^ and a logK of 4.24 ± 0.30. Further analysis resulted in a small yet favourable entropic contribution ΔS of (9.4 ± 4.3) J K^-1^ mol^-1^, a ΔG of (-24.20 ± 1.7) kJ mol^-1^ (**Fig. 3b**), and a K_d_ of (57 ± 17) μM. The latter value is about one magnitude higher than determined by QCM-D. The reason is likely because ITC detects the Gal-1 membrane interaction in bulk under defined concentrations and equilibrium conditions, while QCM-D characterizes the Gal-1 binding to an immobilized 2-dimensional membrane system with reproducible surface coverage by LUVs (see **Table S3** in the **S.I**.). Although the enthalpy involved is of the same magnitude as the one reported for Gal-1 binding to bivalent disaccharides containing terminal galactose moieties, the entropies significantly differ. ^50, 51^ The entropy increase, related to the release of the water molecules from the Gal-1 CRD cavity upon binding to galactose of GM_1_ or bivalent disaccharides, is comparable. ^52^ However, disaccharides diffuse in the solution ^50, 51^ while GM_1_ is embedded in the lipid membrane, which gives a restricted number of starting configurations. Subsequently, a more constrained system undergoes a less significant loss in the degrees of freedom upon binding. On the other hand, the enthalpy is in the same range, pointing towards the involvement of the terminal galactose moiety of the GM_1_ in the interaction. The careful analysis of the ITC data suggests a 1:1 Gal-1:GM_1_ binding stoichiometry, which can be interpreted as an argument against the suggested GM_1_ cross-linking by Gal-1 homodimer. That cross-linking is suggested to lead to the formation of a “glycan-galectin aggregation”. ^53^ Although that concept has gained wide popularity, only very few studies show indirect proof of aggregate formation ^36^, specifically for Gal-1 homodimer, a proof for GM_1_ cross-linking is missing.

In the context of GM_1_ cross-linking by Gal-1 it has to be considered that Gal-1 is mainly monomeric in solution at physiological concentrations. ^54^ This finding agrees with the K_d_s reported in the literature in the micromolar range (2.5 ^55^ or 7 μM ^56-58^, respectively) and with our Fluorescence Correlation Spectroscopy experiments (see **S.I**.). Taking this into consideration, in the used concentration range from 2 to 10 μM, we expect to have between 80% and 40% of Gal-1 monomer in solution, depending on which K_d_ is used (for further details see **S.I**. and **Fig. S4**). Similarly, some cell types express galectins at micromolar cytosolic concentrations, ^54, 59^ Thus the majority of Gal-1 might be present as monomers not favoring cross-linking. However, cross-linking might still occur due to local accumulation of Gal-1 at the cell membrane upon specific cellular stimuli leading to significantly higher local Gal-1 concentrations than used in our study (10 μM). Since galectins seem to possess more affinity for glycoproteins rather than glycolipids, ^54^ Gal- 1/glycoproteins binding could cause the increase of its local concentration, which might favor the subsequent binding of the Gal-1 homodimer to GM_1_. One might speculate that the elevated concentration of membrane bound Gal-1 might in turn then lead to cross-linking, stabilizing the transient GM_1_ nanodomains, ^40, 41^ and by that the formation of stable “GM_1_-Gal-1 lattices”. At this point of the discussion we shall mention further aspects reported in the literature relevant to the molecular mechanisms involved in that cross-linking: ligand binding to Gal-1 increases its affinity to form dimers, ^60^ and a negative cooperativity was found for the carbohydrate-binding to the dimeric form. ^61, 62^

All-atom MD simulations offer further insight into the stability of Gal-1 homodimer and interactions of the homodimeric Gal-1 with different lipid species, which are not easily accessible by experiments. To address the formation of Gal-1 homodimer, claimed to be a prerequisite for cross-linking of GM_1_ lipids or GM_1_- associated integrin, ^27^ we assessed the stability of Gal-1 homodimer in water by calculating the distance of the center of mass (COM) of the two monomers and by analyzing the secondary structure content (**Fig. S7**). **Fig. S6** shows the time evolution of the COM distance between Gal-1 monomers in the homodimer. This distance remains constant, with an average of 2.89 ± 0.02 nm, suggesting that the homodimeric Gal- 1 is stable throughout the entire simulation (three repeats, each 1 μs long). To assess the changes in the secondary structure of the protein, we plot the time dependence of the secondary structure content for all repeats. As shown in **Fig. S7**, there are no substantial changes in the secondary structure content of Gal- 1 homodimer, again suggesting high dimer stability.

To gain further insight into the stability of the dimeric interface, we pulled Gal-1 monomers apart and calculated the free energy needed to dissociate the homodimer using the umbrella sampling technique. ^63, 64^ **Fig. 3c** demonstrates that the homodimeric structure of Gal-1 is preferred and that the free energy difference between the dimeric and monomeric states is about 140 kJ/mol. Notably, the conditions in the MD simulations relate to a bulk concentration in the sub-mM range. Experiments yield K_d_s in the range of a few μM for the Gal-1 monomer-dimer equilibrium ^55-58^ and thus confirm a stable homodimer at sub-mM concentrations, as shown in the simulations.

Subsequently, we explored the interactions of Gal-1 homodimer with model lipid membranes by assessing the Gal-1average interaction time and binding probability to different lipid species (**Fig. S8** and **Fig3d**, respectively). The Gal-1 dimer was initially placed about 2 nm above the bilayer surface and allowed to move freely in the simulation box. Five different lipid bilayer compositions have been used: POPC, POPC:GM_1_ (96/4), POPC:GM_1_ (90/10), POPC: (96/4), and POPC:GD_1_a (90/10), (see **Table S2** for details). The binding probabilities of Gal-1 dimer to different lipid species (shown in **Fig. 3d**) have been calculated by counting the number of simulation frames where Gal-1 dimer was in contact (within the distance of 0.6 nm) with the given lipid type. Our calculation unveiled intriguing patterns in the interaction dynamics between GM_1_ and the Gal-1 dimer. The binding probability experiences a notable upswing as the concentration of GM_1_ increases. Conversely, the binding probability to GD_1_a exhibits a diminishing trend with increasing GD_1_a concentration. A comparative analysis with a pure POPC bilayer reveals that both POPC:GM_1_ and POPC:GD_1_a systems demonstrate somewhat elevated binding probabilities to the Gal-1 dimer (except for the system containing 10 mol% of GD_1_a, where overall Gal-1 homodimer binding is slightly smaller as compared to pure POPC bilayer). It is essential to underscore that the binding probability to POPC is not negligible either, indicating the capacity of the Gal-1 dimer for unspecific binding to lipid bilayers, even in the absence of ganglioside lipids. In summary, the trends observed in the MD simulations support our experimental view: Gal-1 binds specifically to GM_1_, but not to GD_1_a, and sub-mM concentrations of Gal-1 might favor the proposed GM_1_ cross-linking by Gal-1 homodimers.

## Conclusions

Synergistically with the binding to glycoproteins, the conversion of GD_1_a to GM_1_ supposedly triggers the binding of Gal-1 to the outer layer of the plasma membrane. This first concept is based on the elevated binding affinity of Gal-1 towards GM_1_ over GD1a when embedded in membranes, an assumption lacking experimental proof. In summary, our study unambiguously demonstrates that when probing a wide range of concentrations (i.e., 30 nM to 10 μM) Gal-1 binds indeed specifically to GM_1_, and not to GD_1_a when embedded in membranes at ganglioside concentrations of 4% with apparent dissociation constants of (4 ± 1) μM and (57 ± 17) μM for binding to immobilised and free-diffusing vesicles, respectively.

The second concept regarding the molecular mechanisms involved in the biological function of Gal-1 is the hypothesis of cross-linking of GM_1_’s by homobivalent Gal-1. The analysis of the ITC experiments indicates a 1:1 binding stoichiometry when using Gal-1 concentrations in the μM range. Considering that both, the K_d_s for the Gal-1/GM_1_ interaction and for the Gal-1/Gal-1 dimerization ^55-58^ are in the μM range, the investigated systems using μM concentrations must be in a dynamic equilibrium. Our results imply that forming such a “GM_1_-Gal-1 lattice” ^53^ will certainly need higher than local micromolar Gal-1 concentrations.

This calls for additional mechanisms driving both the Gal-1/Gal-1 dimerization and the GM_1_/Gal-1 equilibrium towards the membrane bound homobivalent Gal-1. *In vivo* this can be achieved by galectin accumulation in extracellular structures rich in galactoside residues, such as extracellular matrix or plasma membrane-proximal glycocalyx as shown before. ^37, 65^ Similar trapping in glycostructures and deposition to surface receptors was described for secreted soluble signaling molecules, such as growth factors. ^66^

## Acknowledgments

The authors acknowledge Annalinda Contino (University of Catania) and Manuel Prieto (University of Lisboa) for their expertise and fruitful discussions. We also thank Giovanni Occhino for his technical assistance, Marco Mauro for his technical support with the QCM-D, Francesco Attanasio (CNR, Catania), Carola Rando and Vladimír Šindelář (Masarykovo University, Brno), Jitka Holková (CEITEC, Brno) for their support with the ITC measurements. We acknowledge CIISB, Instruct-CZ Centre of Instruct-ERIC EU consortium, funded by MEYS CR infrastructure project LM2023042 and European Regional Development Fund-Project „UP CIISB” (No. CZ.02.1.01/0.0/0.0/18_046/0015974) for the financial support of the measurements at the CF Biomolecular Interactions and Crystallography. I.V. and W.K. have been supported by the Academy of Finland (projects 331349, 336234, 346135), the Sigrid Juselius Foundation, Helsinki Institute of Life Science (HiLIFE) Fellow Program, the Human Frontier Science Program (RGP0059/2019), the Lundbeck Foundation, and DFG (SFB/TRR 83). We acknowledge the computing resources provided by the CSC – IT Center for Science Ltd. (Espoo, Finland) and LUMI supercomputer, owned by the EuroHPC Joint Undertaking, hosted by CSC and the LUMI Consortium.G.N. and G.M. thank PIAno di InCEntivi per la RIcerca di Ateneo -PIA.CE.RI. 2020/2022 - linea 2 - Progetto di ricerca intradipartimentale Phototeranostic and Microvescicle Recognition Nanostructures under electromagnetic Activation for financial support. The authors acknowledge the assistance provided by the Advanced Multiscale Materials for Key Enabling Technologies project supported by the Ministry of Education, Youth, and Sports of the Czech Republic. Project No. CZ.02.01.01/00/22_008/0004558, Co-funded by the European Union.

