## Supplemental Material for "Unraveling the GM_1_ specificity of Galectin-1 binding to lipid membranes"

**Supporting Information**

Table of content

#### Experimental Procedures

##### Materials

1-palmitoyl-2-oleoyl-sn-glycero-3-phosphocholine (POPC), Ganglioside GM_1_ (Ovine Brain), 1-palmitoyl-2-oleoyl-sn-glycero-3-phosphoethanolamine-N-(capbiotinyl) (DOPE-cap-biotin) and 1,2-dioleoyl-sn-glycero-3-phosphoethanolamine-N-(lissamine rhodamine B sulfonyl) (ammonium salt) were purchased from Avanti Polar Lipid (Alabaster, AL, USA). Ganglioside GD_1_a disodium salt (bovine brain) was purchased from Enzo LifeScience (Farmingdale, NY, USA). Chloroform (containing amylenes as a stabilizer, ACS reagent, ≥99.8%), methanol (ACS reagent, ≥99.8%), Phosphate Buffered Saline (PBS) (tablets),4-(2-Hydroxyethyl)-piperazine-1-ethanesulfonic acid sodium salt (HEPES), sodium chloride (NaCl), Hydrogen Peroxide Solution (30%), Ammonia (25%), sucrose, Bovine Serum Albumin (BSA) were acquired from SigmaAldrich (St.Louis, MO) (all 99% purity, unless otherwise stated). DOPE (1,2-Dioleoyl-sn-glycero-3-phosphoethanolamine) labeled with Atto-633 was provided by Atto-Tec GmbH (Siegen, Germany). TAMRA maleimide, 6-isomer was purchased from Lumiprobe (Hannover, Germany). Terrific Broth medium (TB) (Roth, Karlsruhe, Germany), Luria Broth medium and Sodium bicarbonate were purchased from Roth (Karlsruhe, Germany), while PD10 column from Cytiva, Merck (Darmstadt, Germany). Biotinylated bovine serum albumin (biotin-BSA) and Cholera Toxin Subunit B (Recombinant), Alexa Fluor™ 488 Conjugate were purchased from Thermo Fisher (Waltham, MA USA) and streptavidin from IBA Lifesciences (Göttingen, Germany).

##### Recombinant protein expression and purification

For recombinant protein expression of the wild type human Gal-1, *Escherichia coli* BL21 (DE3) pLysS cells (Promega) were transformed with the pGEMEX-Gal-1 plasmid. Transformed bacteria were grown for 16 h at 37 °C in Luria Broth medium (Roth, Karlsruhe, Germany) containing the appropriate selection antibiotic. For protein expression medium was changed to Terrific Broth medium (TB) (Roth). After initial growth for 2 – 3 h at 37 °C in TB medium up to an OD600nm of 0.6 – 0.8, gene expression was induced using 100 µM β-1-thio-D-galactopyranoside (IPTG), and bacteria were cultured at 37 °C for additional 16 h. Cells were harvested, washed and bacteria pellets were frozen for 2 h at -20 °C before lysed by sonication at 4 °C. The protein was purified from the bacterial extracts after lysis by affinity chromatography on lactosylated Sepharose 4B. The lactosylated Sepharose 4B column bound protein was labeled directly with a four-molar excess of maleimide-TAMRA (TMR) dye (Lumiprobe, Hanover, Germany) followed by extensive washing steps before the elution. Therefore, the buffer was changed to labeling buffer (0.1 M sodium bicarbonate pH=8.3) and labeling occurred according manufactural instructions rotating overnight at 4 °C in the dark. After labeling, free dye was washed out of the column with 20 mM PBS pH=7.2 buffer and active protein was eluted with 50 mM lactose in 20 mM PBS pH=7.2 followed by buffer exchange to 10 mM PBS pH=7.2 via PD10 column (Cytiva, Merck Darmstadt, Germany) to remove lactose. Purity was ascertained by one- dimensional gel electrophoresis under denaturing conditions.

##### Model membranes preparation

Briefly, the desired lipid compositions vesicles, Large Unilamellar Vesicles (LUVs) or Giant Unilamellar Vesicles (GUVs) were obtained from the chloroform stock solutions using extrusion or electroformation methods. ^1-3^ 1% of DOPE-Atto633 probe or 1% of DPPE-cap-biotin were added to the chloroform solution when needed. The mixture was dried under nitrogen flow in a test tube (LUVs) or spread onto two pre-ozonized and pre-heated titanium plates (GUVs), in both cases further evaporated under vacuum for at least 3 h. The dry lipid films were hydrated and vortexed in a buffer solution (10 mM PBS, 137 mM NaCl, 0.27 mM KCl, pH=7.4) for LUVs preparation. Whereas the lipid-coated plates were assembled and filled with sucrose in water (107 mOsm kg^-1^) and then placed on a heating plate at approximately 45 °C. Before the LUVs extrusion, seven cycles of freeze and thaw were performed, using liquid nitrogen and water bath, set above the solid-liquid transition temperature characteristic of the lipid used (42-45 °C). The dispersion was later extruded (101 times at 45 °C) through polycarbonate filters (pore diameter 100 nm, Nuclepore, Pleasanton, CA) mounted in a mini-extruder (Avanti Polar Lipid, Birmingham, England) fitted with two 0.5 mL Hamilton syringes (Hamilton, Reno, NV). For GUVs preparation, the voltage was increased stepwise from 0.250 V to 3.5 V (peak-to-peak voltage) for 65 min and then kept at 3.5 V for 1 h. In the final step, the frequency was decreased stepwise from 10 to 4 Hz and the voltage kept to 3.5 V for 1h to detach the formed liposomes. The temperature was kept at 45 °C. The obtained GUVs dispersions were later diluted with buffer of the same osmolarity (10 mM HEPES, 50 mM NaCl, pH=7.4, 107 mOsm kg^-1^).

**Table S1.** Lipid compositions used in this study.

| Name | POPC | GM_1_ | GD_1_a |
| --- | --- | --- | --- |
| POPC | 100 | - | - |
| POPC:GM_1_ | 96 | 4 | - |
| POPC:GD_1_a | 96 | - | 4 |

##### Fluorescence Confocal Microscopy and Fluorescence Correlation Spectroscopy

Before the CF measurements, the µ-Slide 8 well-ibidi chambers (Gräfelfing, Germany) chamber with glass bottom was coated with 200 μL of BSA-biot 0.1 mg/ml, 200 μL of streptavidin (2 μg/ml), waiting 30 min each step and washing all the wells with mQ water after each step of coating. In each well, we added 80 μL of GUVs (HEPES 10 mM, NaCl 50 mM, pH=7.4, 107 mOsm kg^-1^) and after letting the GUVs attach to the coated bottom for 30 min, 30 nM of Gal-1/TMR was added and diluted to reach a total volume of 450 μL. Prior to each measurement, the background, i.e. the unlabeled GUVs, was checked. For Fluorescence Correlation Spectroscopy (FCS) experiments, to avoid Gal-1/TMR to adsorb on the glass bottom, we coated the chambers using only BSA. Image acquisition and FCS measurements were performed on a home-built confocal microscope. Pulsed diode laser (532 nm) was used at 25 MHz repetition rate. A quad-band dichroic mirror (375/470/532/640) was used to up-reflect the light onto a water immersion objective (60x, NA 1.2). Furthermore, a 570-620 nm bandpass filter was employed. For imaging acquisition, the power of the laser was kept below 5 μW and each image was recorded at a different resolution depending on the size of the single GUV, scanning in monodirectional mode. The images were acquired after 1 h from the addition of the protein to the GUVs chamber. The experiments were performed twice, with two different sets of electroformed GUVs (two biological replicates, overall 10 GUVs were analyzed). For FCS the power of the laser was set to 10 μW (measured at the end of the fiber) and each point was acquired for 2 min.

##### Förster Resonance Energy Transfer (FRET)

Time-resolved measurements were performed by using a modular FluoTime300 Spectrofluorometer (PicoQuant GmbH, Berlin, Germany) using Time-correlated single photon counting (TCSPC). The setup has been equipped with green excitation laser (PicoQuant LDH-D-TA 532, pulse width less than 100 ps, 80 MHz maximum repetition rate, emission peaking at 531 nm) and a HPMA-06 hybrid photomultiplier tube. The overall instrument response function width was around 120 ps FWHM. An emission long-pass cutoff filter (540 nm) was used to eliminate the scattered excitation light. The sample was measured at 578 nm emission wavelength. The monochromator slit-width was adjusted according to the protein emission intensity and then kept constant during the whole titration. Titrations were performed by adding different volumes of red-labelled LUVs of different compositions (4 mM) to 0.5 μM of Gal-1/TMR into an Ultra-Micro Cell cuvette (Hellma, Merck KGaA, Darmstadt, Germany). After 10 min from each addition a decay curve was recorded with the setup described above. The so-obtained set of decays were analyzed globally, using iterative reconvolution fitting of a three-exponential function using EasyTau 2 Software (PicoQuant GmbH, Berlin, Germany). The Amplitude Weighted Average Lifetime has been calculated and plotted (together with its standard deviation, n=5, four different extrusions for POPC and POPC:GM1 data, n=2, two different extrusions for POPC:GD1a data).

##### Quartz Crystal Microbalance with Dissipation monitoring (QCM-D)

The measurements were conducted using a Quartz Crystal Microbalance with Dissipation monitoring system, equipped with a quartz crystal of 5 MHz (diameter 14 mm) (Novatech Srl, Pompei, Italy). Before each measurement, the sensor had to be properly cleaned and activated as previously described.^4, 5^ Briefly, a solution 5:1:1 of distilled water, ammonia solution (25%) and hydrogen peroxide (30%) was used to treat the sensor at 70 °C for 20 min. Following, the sensor was rinsed with water and dried under nitrogen flow, then exposed under a UV lamp for 10 min, twice. Before performing each measurement, the sensor was calibrated and then both the frequency and dissipation changes at first overtone were monitored previously in the air, then following the injection of the buffer. Once the frequency and the dissipation were stable, the 1 mM of lipid dispersion of the desired composition was injected into the chamber until further stabilization. In the case of POPC:GM_1_ and POPC:GD_1_a, the observed frequency shifts were smaller than the ones related to POPC depositions, due to an incomplete coverage of sensor. Therefore, after the deposition of the vesicles containing gangliosides, the inert dispersion of POPC was perfused until saturation of the sensor, corresponding to the stabilization of the frequency value. Once the frequency reached a constant value, the Gal-1 solution (10 mM PBS, 137 mM NaCl, 0.27 mM KCl, pH=7.4) was introduced into the chamber, succeeded once again by the buffer. Each injection step was followed again by the buffer perfusion. A flow rate of 20 µL/min was used and kept constant during the whole experiment.

##### Isothermal Titration Calorimetry (ITC)

The experiments were performed by using an ITC Nano Active Control Calorimeter (TA Instruments) and Auto-iTC200 (Malvern). The first one was equipped with a gold cell (volume of 988 µL, provided by the manufacturer), whereas the second one was equipped with a coin-shaped Hastelloy cell (200 µL). Three syringes, 40, 100 and 250 μL volume equipped with a shaker, have been used to perform the titrations. Each solution used has been previously degassed for 20 min. This time was optimized to minimize the variations in the concentration of the prepared solutions. The reliability of the results obtained with the ITC Nano Active Control Calorimeter was further strengthened by the precision and accuracy of the calorimetric apparatus, thoroughly tested by using a chemical calibration procedure ^6^ rather than the simple electrical calibration recommended by the ITC manufacturers. ^7, 8^ The concentrations used in the experiments are within the range 2-10 µM and 1.5 mM for the Gal-1 and the LUVs, respectively. All the experiments were carried out in overfill mode at 25 °C, in a temperature-controlled room (25.0 ± 0.4 °C), and under stirring (750 rpm).

##### Molecular Dynamics (MD) Simulations Setup

Molecular dynamics (MD) simulations were carried out on five protein–lipid bilayer systems, whose detailed molecular compositions are given in Table 2. In each of these systems, the homodimeric Gal-1 (PDB ID: 1GZW) was placed at least 2 nm above the surface of the lipid bilayer in three random orientations. An appropriate number of water molecules was added to solvate the system fully. We added 0.15 M KCl and inserted additional potassium ions to neutralize the net charge of the system. All molecules in the system were described using the CHARMM36m force field. ^9^ The resulting 15 MD simulations were energy minimized and 1 μs long simulations were carried out with a time step of 2 fs. The first 500 ns of the simulation time were considered as equilibration time and excluded from subsequent analyses. The periodic boundary conditions were used in all three dimensions. The simulations were performed in the NpT ensemble with a temperature of 298 K kept using the Nosé-Hoover thermostat ^10, 11^ and a pressure of 1 atm kept by Parrinello-Rahman barostat. ^12^ The coupling constants for temperature and pressure were 1 ps and 5 ps, respectively. Temperature coupling for protein, lipid bilayer, and solvent (water and ions) were independent. The pressure in the membrane plane (*xy* plane) was maintained independently from pressure along bilayer normal (semi-isotropic pressure coupling). All simulations were performed using GROMACS 2020.5 software. ^13^

The calculation of the potential of mean force (PMF) between the Gal-1 monomers in water was carried out using the umbrella sampling technique. ^14, 15^ The Gal-1 homodimer was placed in the middle of the cubic simulation box. The system was solvated and potassium ions were added to neutralize the net charge of the system. Before the beginning of the pulling simulation, the system was energy minimized and equilibrated for 200 ns. Thirty-four windows with 0.1 nm spacing were constructed by pulling apart the centers of mass of Gal-1 monomers. The pulling rate of 0.2 nm/ns with 2000 kJ mol^-1^ nm^-2^ force constant was used. Each window was simulated for 200 ns with the first 100 ns considered as equilibration time and removed from the free energy calculations. The PMF was calculated using the weighted histogram analysis method implemented in GROMACS. ^16^ Errors were estimated using the Bayesian bootstrapping method. ^17^

**Table S2**. Molecular compositions of the systems used in the MD simulations.

| Label | Gal-1 homo-dimer | POPC | GM_1_ | GD_1_a | K^+^ | Cl^-^ | water | Number of replicates | Simulation time per replicate (µs) |
| --- | --- | --- | --- | --- | --- | --- | --- | --- | --- |
| POPC | 1 | 256 | - | - | 28 | 22 | 32800 | 3 | 1 |
| POPC:GM_1_ (4 mol%) | 1 | 246 | 10 | - | 68 | 52 | 41541 | 3 | 1 |
| POPC:GM_1_ (10 mol%)* | 1 | 230 | 26 | - | 80 | 48 | 40254 | 3 | 1 |
| POPC:GD_1_a (4 mol%) | 1 | 246 | - | 10 | 83 | 57 | 43150 | 3 | 1 |
| POPC:GD_1_a (10 mol%)* | 1 | 230 | - | 26 | 114 | 56 | 42846 | 3 | 1 |

* A 10 mol% GM_1_ has been used in the MD simulations to enhance the sampling.

### Results and Discussion

#### Characterization of Gal-1/TMR

##### Choice of the FRET pair and calculation of the Förster Radius

The lifetime of Gal-1/TMR as a function of different concentrations of vesicles has been monitored to study their interaction. The labeled protein has been used as a FRET donor, whose fluorescence emission spectrum is reported in **fig. S1** (magenta curve). Considering the significant overlap with its UV-Vis spectrum (**fig. S1**, cyan curve), qualitatively highlighted in the **fig. S1** as a pale purple area, DOPE-Atto 633 (1% of the total lipid concentration) has been used as the FRET acceptor. This lipid conjugated dye is routinely employed in a large variety of fluorescence studies due to the feature of being easily intercalated into the lipid bilayer. ^18^

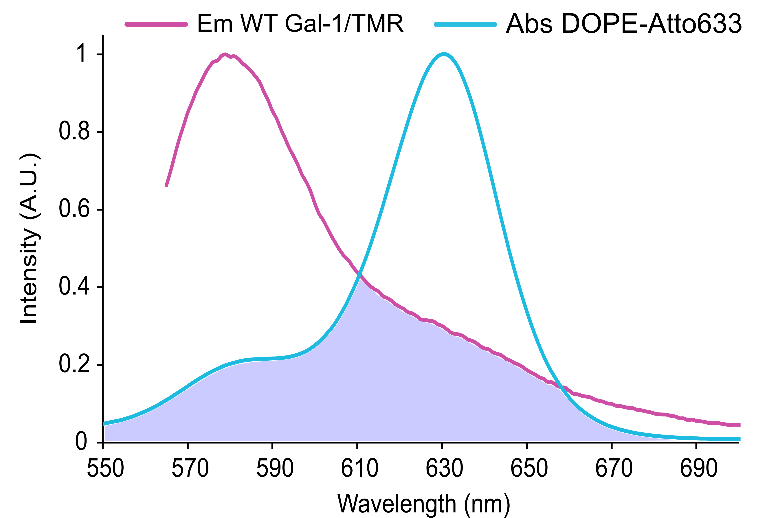

**Figure S1**. FRET pair used for the fluorescence measurements. The magenta spectrum shows the emission of Gal-1/TMR (𝛌 _max_ = 580 nm), whereas the light blue curve shows the absorption of the DOPE-Atto633 (𝛌 _max_ = 630 nm). In pale purple the spectral overlap between the two.

The Förster Radius of the above-described FRET pair was calculated by using [1]. ^19^

${R_{0}={0.211(\kappa}^{2}n^{-4}Q_{D}J_{(\lambda)})}^{1/6}$ [1]

Where $\kappa^{2}$ is a factor considering the relative orientation of the FRET pair’s transition dipoles, usually approximated to 2/3, ^19^ $n$ is the refractive index of the solvent and $Q_{D}$ is the quantum yield of the donor. $J_{(\lambda)}$is the overlap integrand and it is defined as:

$$J_{(\lambda)}= \int_{0}^{\infty} F_{D(\lambda)}\varepsilon_{A(\lambda)}\lambda^{4}d\lambda[2]$$

Where $F_{D(\lambda)}$ is the corrected fluorescence intensity of the donor with the total intensity, between $\lambda$ and $\lambda+ \Delta\lambda$, and the $\varepsilon_{A(\lambda)}$ is the extinction coefficient of the acceptor as a function of $\lambda$.
For the FRET pair used in our study, (i.e. TMR and Atto-633 respectively conjugated to the protein and the lipid in the membrane, **fig.S1**) we determined a Förster radius of 45 Å.

##### Calculation of Degree of Labeling (DOL)

Gal-1 was labelled with TMR-maleimide following the procedure reported in **Materials and Methods**. TMR-maleimide has been chosen because it selectively reacts via cysteine, also preventing the protein oxidation which would affect its activity. The Gal-1 possesses six cysteines per monomer, i.e., Cys2, Cys16, Cys42, Cys60, Cys88 and Cys130, two of them (C2 and C135) are the most exposed to the solvent, ^20^ thus the most reactive, and also the one responsible for the tertiary and quaternary structures. ^21^ The DOL is defined as the ratio between the moles of the dye per moles of Gal-1, and it was calculated using UV-Vis spectroscopy. The spectrum was acquired at room temperature using a Shimadzu Europe UV-2600 spectrophotometer in the range of 200–700 nm. Briefly, the concentration of TMR was measured by monitoring the absorbance at 552 nm and the concentration of Gal-1/TMR considering the absorbance at 280 nm, to which the contribution of the dye has been subtracted, knowing from other experiments the extinction coefficient of the dye at 280 nm and the respective concentrations (**fig. S2**).

The UV-Vis spectrum at 280 nm is attributed to two contributions, i.e., the TMR and the Gal-1, as expressed in the **eq. 3**.

$$A_{280}=A_{280 TMR}+ A_{280 Gal-1}\left[ 3 \right]$$

$$A_{280}=C_{TMR}\varepsilon_{280 TMR}+C_{Gal-1} \varepsilon_{280 Gal-1}[4]$$

$$C_{Gal-1}=\frac{A_{280}-C_{TMR}\varepsilon_{280 TMR}}{\varepsilon_{280 Gal-1}}[5]$$

where C is the concentration, A is the measured absorbance and ε is the molar absorptivity in M^-1^ cm^-1^. The concentration of the dye (C_TMR_) was obtained using the A_552nm_ of the measured spectrum (**Fig. S3A**) due to the TMR only. The ε_552_ was calculated by measuring three UV-vis spectra at three different concentrations of the TMR-maleimide in mQ water. The slopes of the three different linear curves are the ε at 552 nm, 515 nm and 280 nm (**Fig S2**).

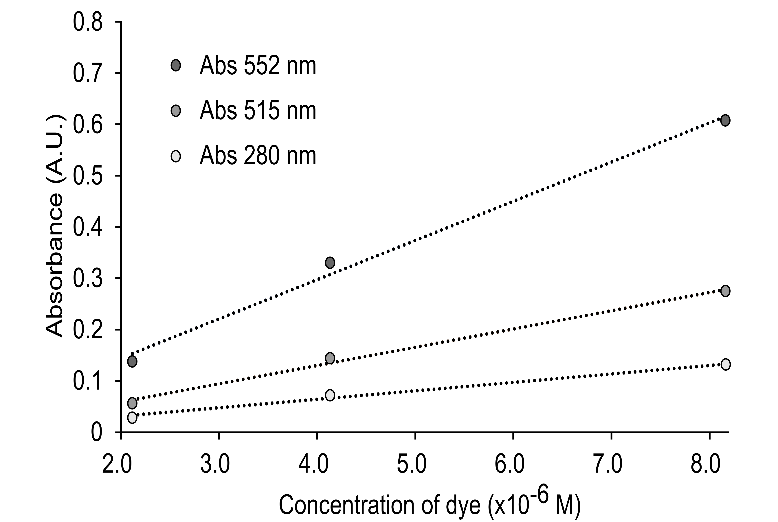
 **Fig. S2.** Absorbance of TMR-maleimide in mQ water as a function of the concentration. Three different wavelengths have been plotted. The R-square are 0.9927, 0.9042, and 0.9884, respectively, for dark grey, grey and light grey curves.

Gal-1’s ε^1%^_1 cm_ is 5.4, as stated elsewhere. ^22^ We used this value upon conversion into the molar absorptivity expressed in mol^-1^cm^-1^. Using the **eq. 3**, the DOL resulted in 0.6.

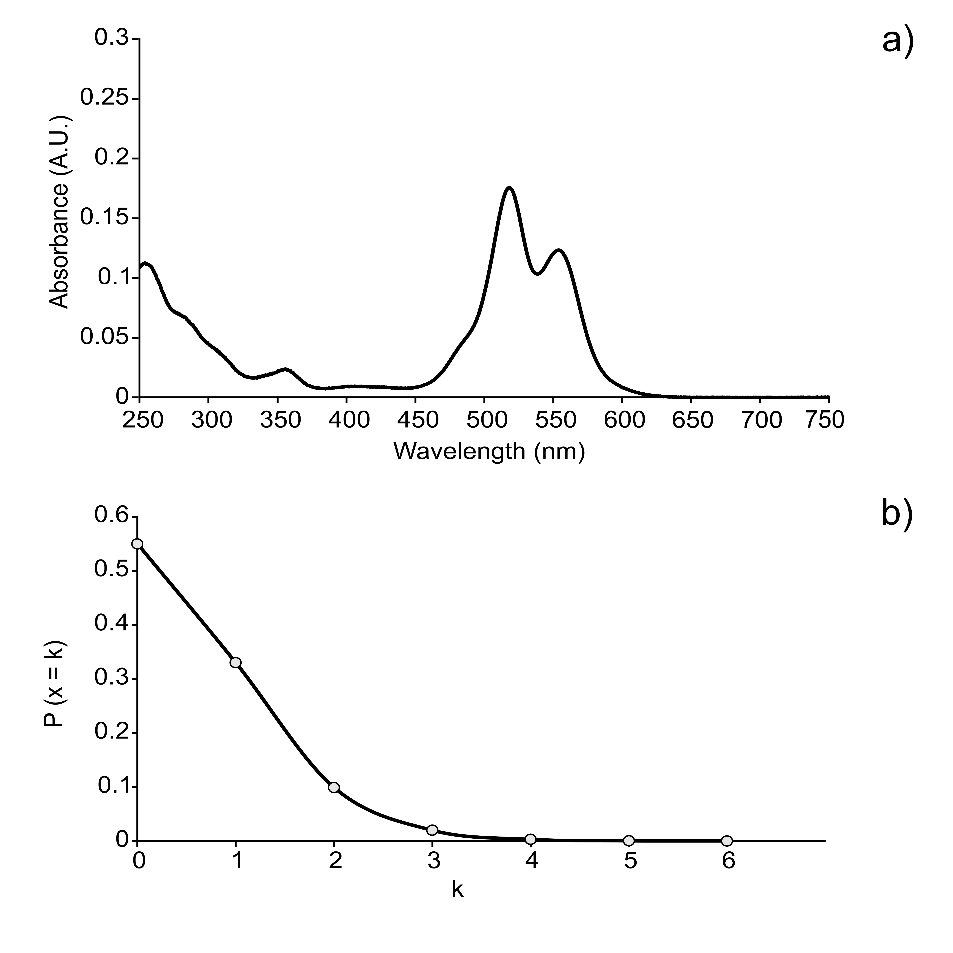

**Figure S3.** A) UV-Vis spectrum of Gal-1/TMR. B) Poisson distribution to the corresponding DOL 0.6.

From a photophysical point of view, TMR dye is known for forming of H-dimers under certain conditions. ^23-25^ Gal-1/TMR is characterized by a significant amount of H-dimers, as shown by the UV-vis spectra reported in **Fig. S3a**. Since the characteristic absorption spectrum of TMR shows a unique maximum at around 552 nm and only a barely visible shoulder at around 520 nm, the pronounced maximum at around 520 nm of the Gal-1/TMR indicates that many Gal-1 molecules carry more than one TMR molecule. ^24^ The Poisson distribution corresponding to a DOL of 0.6 is reported in **Figure S3b**. The distribution shows that there is roughly 55% of the protein which did not react with the dye, thus unlabeled, 33% of the population is represented by the 1:1 (protein to dye) species, 10% by the 1:2 species, and 2% of the protein is labelled with 3 dyes. While these features do not shatter the soundness of the FRET titrations (**Fig. 1c, d, e, f**) since the control is always measured, on the other hand, it might be one of the reasons preventing us from obtaining the classical binding curve trend.

##### Fluorescence Correlation Spectroscopy characterization

To characterize Gal-1/TMR we employed FCS. We acquired single-point measurements in solution, each of them for 2 min. The global analysis of multiple measurements was performed using SymPhoTime 64 Software. The autocorrelation curves were fitted keeping the same boundaries (0.002-1000 ms), choosing a 3D free diffusion model, single species described in the eq. [6] as follows:

$G_{\left( t \right)}=\frac{1}{N}\frac{1}{1+ \left( \frac{t}{\tau} \right)}\sqrt{\left( \frac{1}{1+\left( \frac{t}{\tau} \right)\kappa^{2}} \right)}$ [6]

where *N* is the number of independently diffusing species within the confocal volume, *τ* is the mean diffusion time, and *κ* is the structural parameter describing the shape of the confocal volume. ^26-29^ Size of the confocal volume and *κ* were calculated for aqueous solution of rhodamine-B or the Alexa-532 as a standard (*D* = 427 ± 4 µm^2^/s and 396 ± 10 µm^2^/s at 298.15 K and 295.65 K, respectively). ^30, 31^ The mean diffusion time of the free Gal-1/TMR determined this way was *τ* = 0.138 ± 0.003 ms, and from that we calculated the diffusion coefficient (*D*) using the eq. [7]

$D= \frac{{\omega_{0}}^{2}}{4\tau}$ [7]

where *ω*_0_ is the waist of the confocal volume calculated using rhodamine-B or Alexa-532, as described above. The volume of the confocal volume was (0.77 ± 0.04) fL. This procedure resulted in a *D* = 135 ± 2 μm^2^·s^−1^ (n=26). From the comparison with the values for the diffusion constants reported in ^32^ of Gal-1 monomer and homodimer (i.e., 130 and 105 μm^2^·s^−1^, respectively ), we conclude that Gal-1/TMR is a monomer at 50-100 nM which is in line with literature reported equilibrium constants. ^33-36^

#### Equilibrium dimer-monomer of Gal-1

To evaluate the concentration of dimeric and monomeric species present in solution at different protein concentrations adopted in this study, the equilibrium [8] and the mass balance equation [10] were used.

GG ⇌ G + G [8]

Where GG is the Gal-1 homodimer and G is the Gal-1 in the monomeric form. This equilibrium is associated with a dimerization constant (K) as follows:

$K=\frac{{[G]}^{2}}{[GG]}$ [9]

Finally, according to the mass balance law:

$C_{(Gal-1)}$ = $[G]$+ $[GG]$ [10]

Where $C_{(Gal-1)}$ is the analytical concentration of the protein. Using eq. [9] and [10], we obtain:

${[G]}^{2}$ + $K[G]$ - $KC_{Gal-1}$= 0 [11]

We used the obtained equation [11] to write a python script. The script enabled us to obtain a distribution diagram for the two species present in the equilibrium under study under the wide concentration range utilized in this work (**Fig. S4**). The script only requires the dimerization constants ^33-36^ and the concentration range as inputs.

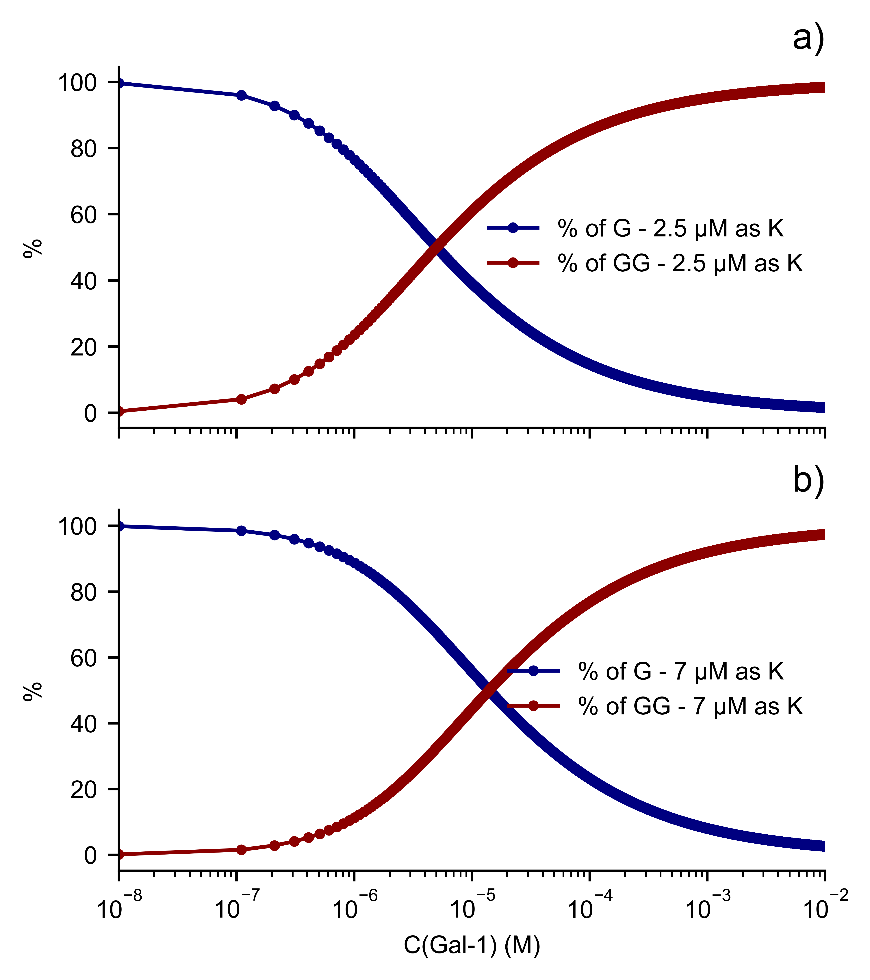

**Figure S4.** Distribution diagram representing the percentage of Gal-1 monomer (G, blue curve) and Gal-1 homodimer (GG, red curve) as a function of Gal-1 concentration utilizing A) a dimerization constant of 2.5 μM ^36^ and B) a dimerization constant of 7 μM. ^33-35^

#### Validation of the FRET methodology

To further validate our FRET methodology reported in the main text (**Fig. 1c, d, e, f**), we took advantage of the high affinity of cholera toxin for GM_1_, which has been extensively studied and reported in previous works. ^37-39^ Cholera toxin has been chosen as a positive control to study the binding to POPC:GM_1_ vesicles, as previously done for Gal-1 and reported in **Fig. 1c, d, e,** and **f**. We performed the same qualitative assay employing commercially available cholera toxin-Alexa488. Its lifetime was monitored as a function of total phospholipids concentration for three different LUVs composition, i.e. POPC (+1% of DOPE-Rhod), POPC:GM_1_ (+1% of DOPE-Rhod) and POPC:GM_1_ without the acceptor. The data are reported in **fig. S4** and clearly show that the effect of the decreasing lifetime as a consequence of the energy transfer can be detected and is not due to any other experimental artifact.

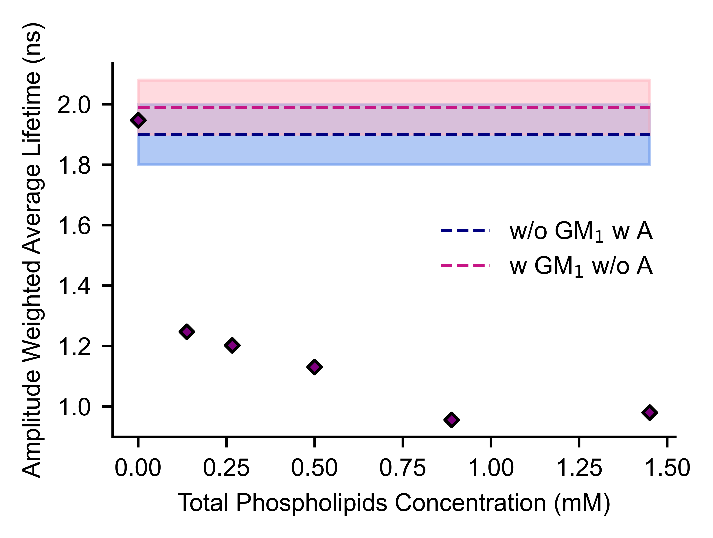

**Figure S5.** Amplitude weighted average lifetime of the cholera toxin-Alexa488 as a function of total phospholipids concentration. Three different compositions were measured. The average of the lifetimes was plotted as a dashed blue line for POPC (+ 1% of DOPE-Rhod as acceptor, named A in the graph). The average of the lifetimes was plotted as a dashed magenta line for POPC:GM1 without acceptor. For POPC:GM1 (96/4) (+1% of DOPE-Rhod) the lifetime was plotted against the concentration of lipid vesicles as purple rhombs.

#### Gal-1 QCM control and fitting of the data to determine the K_d_

We employed QCM-D to both qualitatively show the specificity of Gal-1 for GM_1_ containing membrane and quantitatively determine the apparent K_d_ of the binding to POPC:GM_1_ LUVs deposited onto the gold sensor (**Fig. 2a, c** and **d**). The flow of the experiment is represented in **Fig. 2b**. Briefly, each independent experiment consisted of three steps after treating the sensor, measuring the frequency in air and in the PBS buffer (PBS 10 mM, NaCl 157 mM, KCl 0.27 mM, pH=7.4): I) Injection of the desired amount of vesicles (in huge excess compared to the available surface of the sensor) until stabilization of the frequency, followed by buffer perfusion; II) Injection of inert lipid composition, i.e., POPC, again followed by buffer. This step was introduced to reach the maximum coverage of the sensor and prevent its interaction with the protein; III) perfusion of the desired amount of Gal-1, followed by buffer perfusion after stabilization of the frequency. The frequency shift (Δf) was calculated for each step after the related buffer injection. The washing step with the buffer ensures the exclusion of all the material which is previously deposited of the sensor in a-specific manner. The sensor was treated as described elsewhere ^5^ so that the pre-extruded vesicles would physically adsorb onto the gold sensor without bursting, therefore without forming supported lipid bilayers. However, due to geometric and sterically constrained, this method renders the complete coverage of the sensor a difficult task to achieve. The first experimental precaution was to carry out the second step when injecting compositions different from POPC. The vesicles deposition is rather a stochastic phenomenon, not only based on gravity and diffusion laws, but also affected by electrostatic interactions between vesicles themselves, and with the sensor. The matter complicates when thinking that the size of the vesicles is different and it is a distribution of sizes. All these factors render the maximum frequency variation obtained by injecting vesicles of various compositions different. However, a high degree of coverage is desirable to avoid interferences coming from the direct interaction of the Gal-1 with the naked sensor, especially considering that Gal-1 possesses six cysteine residues per monomer. The derived frequency shift would be summed to the eventual frequency shift caused by the direct interaction with the lipid membranes and the two contributions cannot be separated nor distinguished. This possible artifact would bias the binding detection and the subsequent K_d_ determination. To exclude this scenario in the context of the determination of the binding constant (POPC:GM_1_ only, **Fig. 2d**), we replicated the experiment three times for each concentration, and we focused on the possible correlation between Δf due to the different steps (I, II and III – **Fig. 2b**). The Δf were obtained by subtracting the frequency before the injection to the frequency reached after the washing step performed after each step. The Δf related to the three different steps are reported in **Table S1**.

**Table S3**. Frequency shifts (Δf) of the three different steps in the QCM-D experiments on POPC:GM_1_ (96/4).

| **Gal-1 Concentration (µM)** | **Δf (I step) (Hz)** | **Δf (II step) (Hz)** | **Δf (III step) (Hz)** | **Sum of Δf (I and II steps) (Hz)** |
| --- | --- | --- | --- | --- |
| 0.75 | 211 | 13 | 1 | 224 |
|  | 215 | 17 | 7 | 232 |
|  | 204 | 11 | 8 | 215 |
| 1.25 | 244 | 4 | 9 | 248 |
|  | 227 | 17 | 13 | 244 |
|  | **191** | **6** | **5** | **197** |
|  | 258 | 4 | 13 | 262 |
| 2.50 | 198 | 10 | 12 | 208 |
|  | 246 | 17 | 17 | 263 |
|  | 226 | 17 | 22 | 243 |
| 5.00 | 262 | 7 | 19 | 269 |
|  | **263** | **10** | **16** | **273** |
|  | 238 | 18 | 24 | 256 |
| 10.00 | 209 | 22 | 25 | 231 |
|  | 206 | 12 | 28 | 218 |

The sum of the I and the II steps is directly proportional to the quantity of vesicles deposited onto the sensor prior to the perfusion of Gal-1. As they oscillate between 197 Hz and 273 Hz (bold lines in the table), these values feature a high variability, probably due to the different size distribution of the extruded vesicles (around 120 nm, measured with the Dynamic Light Scattering - data now shown). To help the reader visualize the data, those values were divided into three groups (highlighted in yellow, green and light blue, respectively). The first centered around 210 Hz, the second centered around 235 and the last one centered around 260 Hz. If the highest amount of Gal-1, thus the highest frequency shift, was not due to the binding of the protein with the vesicles but to nonspecific interaction with the gold sensor, the amount of Gal-1 detected on the sensor (with pre-deposited vesicles) would be inversely proportional to the degree of coverage, i.e., the sum of the frequency shifts of the I and the II steps. In other words, the fewer vesicles are on the sensor, the less the frequency, and the more the area exposed of the sensor, the more Gal-1 should be sensed during the third step. This is evidently not our case, as shown by the data reported in the table considering both the collective trend and the trend of the three replicates at the same protein concentration.

This modification of the protocol was needed to avoid potential artifacts stemming from the direct interaction between the protein and the gold sensor. Indeed, this was necessary to differentiate between the two contributions to the frequency shift, i.e., the protein interacting with the lipid vesicles and/or with the sensor (see **S.I., Table S3** to see the data related to step I and II, i.e., vesicles deposition). A schematic drawing of the experiment's flow is depicted in **Fig. 2b**. The data reported in **Table S3** and represented in **Fig. 2d** were used to determine the dissociation constant of the Gal-1 binding to POPC:GM_1_ (96/4) vesicle deposited onto the sensor. The frequency shifts observed upon protein injection (III step) were plotted against the concentrations of Gal-1 used. Those data were fitted using the equation reported in previous work. ^40^

#### Stability of the Gal-1 homodimer

To assess the stability of the Gal-1 homodimer in the MD simulations, the center of mass distance between the Gal-1 monomers in solution has been plotted in **Fig. S6**. Data clearly show that the homodimer is very stable, with an average distance of (2.89 ± 0.02) nm between the monomers.

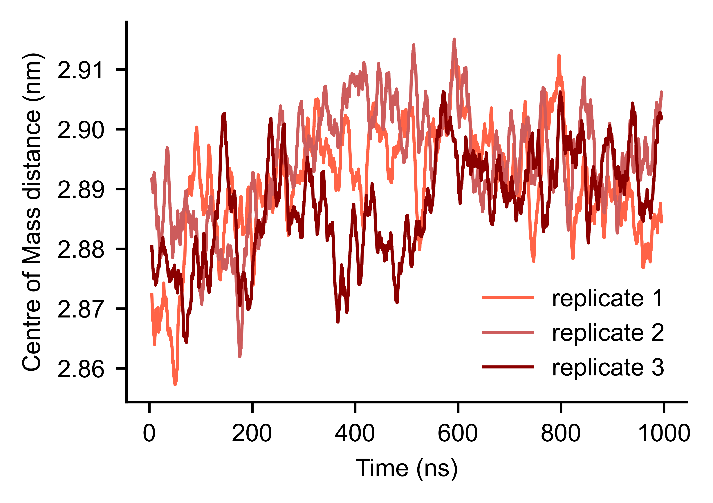

**Figure S6.** Time-dependence of the center of mass distance between Gal-1 monomeric units in the Gal-1 homodimer in solution obtained from the all-atom MD simulations. Data collected from three independent simulations plotted in the three different shades of red.

Additionally, the secondary structure content has been analyzed and depicted in **Fig. S7**. No substantial changes in the secondary structure content are observed suggesting high stability of the Gal-1 homodimer.

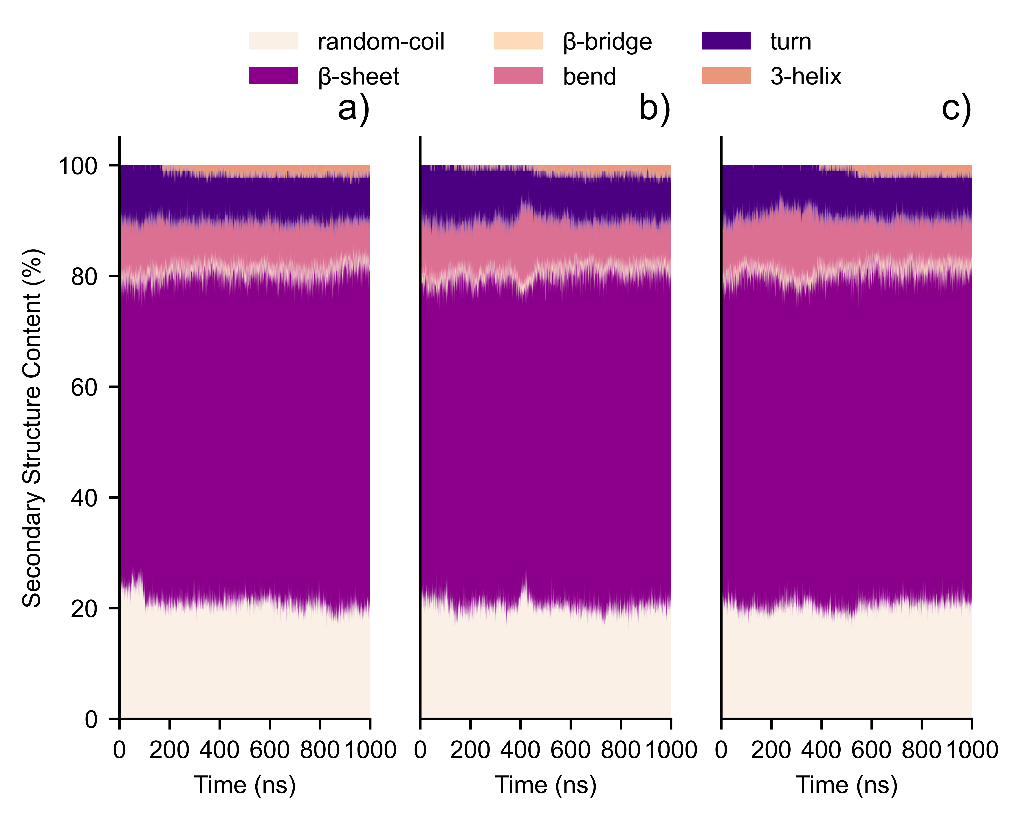

**Figure S7.** Time-dependence of the secondary structure content of the Gal-1 homodimer in solution. Each panel presents the secondary structure content from a single MD simulation. Different types of the secondary structures are color-coded as follows: random coil – linen gray, beta-sheet – dark magenta, beta-bridge - peach, bend – pale violet, turn - purple and 3-helix - dark salmon.

#### Interactions of Gal-1 dimer with lipid bilayers

In addition to binding probabilities showed in the manuscript (see **Fig. 3d**), we calculated the average interaction times between the Gal-1 homodimer and lipid membrane depicted in **Fig. S8**. The average interaction times and standard deviations were calculated from the decay of the time auto-correlation functions by identifying the times when the time auto-correlation function decreased to zero. Results depicted in **Fig. S8** clearly indicate that the average interaction times between the Gal-1 dimer and lipid membrane containing GM_1_ are higher as compared to the interactions with lipid bilayer containing GD_1_a.

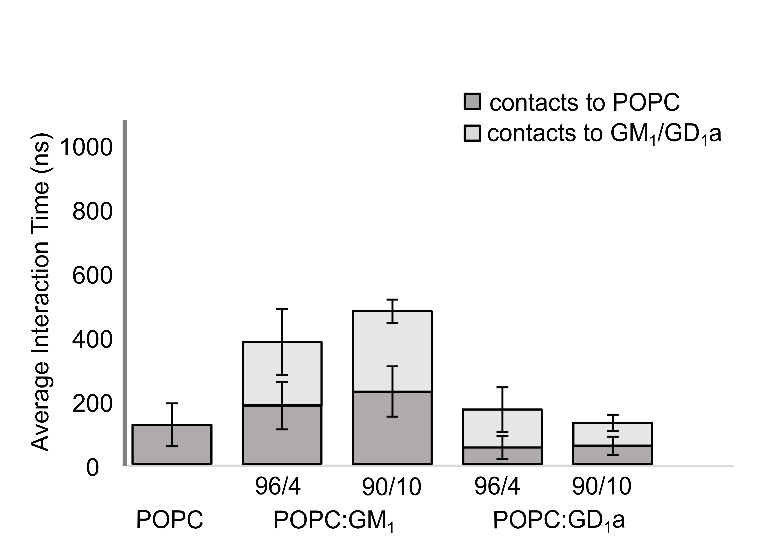

**Figure S8.** Average times and associated standard deviations of the interaction between Gal-1 homodimer binding to five different bilayers, i.e., POPC, POPC:GM_1_ (96/4), POPC:GM_1_ (90/10), POPC:GD_1_a (96/4), POPC:GD_1_a (90/10), represented by the different bars. The bars’ dark and pale grey portions represent the contacts to POPC and GM_1_ or GD_1_a in each different bilayer composition, respectively (see **Table S2** for details).

(32) Göhler, A. Untersuchung Karbohydrat-bindender Proteine mit hoher zeitlicher und

räumlicher Auflösung. University Wuerzburg, 2012.

#### Author Contributions

F.S., G.N., V.Z., and G.M. carried out the experiments. F.S. and G.N. analyzed the experimental data. A.K.L. expressed and purified the protein. J.C.R. measured preliminary data, contributed to the planning and discussion. F.S. prepared the figures. F.S. and G.M. conceived and planned the experiments. F.S., G.N., A.K.L., M.C., G.M., M.H. contributed to the interpretation of the results. P.K. contributed and supervised the technical and instrumental part. P.K., G.M. and M.H. supervised the experiments. W.K. carried out and analyzed the simulations. W.K., I.V. discussed and co-designed the simulations and contributed to the interpretation of the results. F.S., W.K., A.K.L., H.K. and M.H wrote the manuscript. H.J.G. conceived the original idea. A. K. L., H.K, G.M. and M.H. supervised the project. All authors contributed to the discussion and provided critical feedback.
